# Climatic effects on turnover of lowland forest bird communities across a precipitation gradient

**DOI:** 10.1101/251264

**Authors:** Juan Pablo Gomez, José Miguel Ponciano, Scott K. Robinson

## Abstract

One of the main goals of community ecology is to understand the influence of the abiotic environment on the abundance and distribution of species. It has been hypothesized that dry forests are harsher environments than wet forests, which leads to the prediction that environmental filtering should be a more important determinant of patterns of species abundance and composition than in wet forest, where biotic interactions or random assembly should be more important. We attempt to understand the influence of rainfall on the abundance and distribution of bird species along a steep precipitation gradient in an inter-Andean valley in Colombia. We gathered data on species distributions, abundance, morphological traits and phylogenetic relationships to determine the influence of rainfall on the taxonomic, functional and phylogenetic turnover of species along the Magdalena Valley. We demonstrate that there is a strong turnover of community composition at the limit of the dry forest. The taxonomic turnover is steeper than the phylogenetic turnover, suggesting that replacement of closely related species accounts for a disproportionate number of changes along the gradient. We found evidence for environmental filtering in dry forest as species tend to be more tolerant of higher temperature ranges, stronger rainfall seasonality and lower minimum rainfall. On the other hand, wet forest species tend to compete actively for nest space but not for the resources associated with the axes we measured. Our results suggest that rainfall is a strong determinant of community composition when comparing localities above and below the 2400 mm rainfall isocline.

## 1 Introduction

Comparisons of species composition of different communities have long have been used to infer the mechanisms underlying community assembly (Whittaker, 1960). Environmental gradients are particularly useful for such purposes because they potentially allow us to separate the influence of stochastic and niche-based processes (Chase & Myers, 2011; Legendre *et al*., 2005; Tuomisto & Ruokolainen, 2006) and to predict the determinants of ecological communities (Whittaker, 1960; Terborgh, 1977; Jankowski *et al*., 2009, 2013; Condit *et al*., 2002; Swenson *et al*., 2011). In particular, latitudinal and elevational gradients have been studied intensively to help identify the roles of different biotic and abiotic variables in determining community composition (Terborgh, 1977; Jankowski *et al*., 2009, 2013; Swenson *et al*., 2011; Qian & Ricklefs, 2007; Kraft *et al*., 2011; Rodríguez & T Arita, 2004).

Environmental gradients are almost always associated with a change in the harshness of the abiotic environment. At high elevations, for example, low temperatures and high temperature variability are thought to be analogous to the harsher conditions at high latitudes. Such conditions should increase the influence of the abiotic environment on the occurrence and abundance of species (Graham *et al*., 2009; Qian & Ricklefs, 2007, but see Kraft *et al*., 2011). At low elevations and latitudes, where the environment becomes more stable, productivity increases, which also increases the potential for intra- and inter-trophic biotic interactions to determine community composition (Jankowski *et al*., 2012; Martin, 1988; Janzen, 1970). Thus, the turnover of species along elevational and latitudinal gradients is hypothesized to be the result of a change in the relative importance of abiotic and biotic mechanisms that determine community assembly.

Rainfall gradients also potentially vary in climatic stability and harshness. Along the rainfall gradient it is likely that water restricts the distribution of organisms at the dry end and biotic interactions potentially determining community composition in the wet end (Engelbrecht *et al*., 2007; Jabot *et al*., 2008). Plant communities along rainfall gradients, for example, are known to respond dramatically to drought conditions (Condit *et al*., 2002; Engelbrecht *et al*., 2007; Jabot *et al*., 2008). Alternatively, within the same habitat, plants also show niche partitioning, a possible response to competition at different life stages. Pathogens, herbivores, and seed predators also affect plant community composition.

Combining metrics of compositional, functional and phylogenetic beta diversity could increase the power of studies of species turnover on gradients (Stegen & Hurlbert, 2011). Comparisons of functional traits and phylogenetic relationships among species might give additional insights into the mechanisms underlying community composition (McGill *et al*., 2006; Petchey & Gaston, 2006; Graham & Fine, 2008; Bryant *et al*., 2008). The expectations of the functional and phylogenetic turnover differ when the communities are assembled deterministically or stochastically along gradients (Swenson *et al*., 2011; Graham & Fine, 2008). Stochastic mechanisms such as random colonization and extinction predict that while compositional turnover can be high, functional turnover should be similar to that expected by chance (Swenson *et al*., 2011).In contrast, deterministic community assembly predicts high functional turnover among habitat types and low turnover when comparing similar types of habitats.

Both stochastic and deterministic turnover have been documented in plant and animal communities at different geographic and environmental scales (Hubbell, 2001; Gomez *et al*., 2010; Qian & Ricklefs, 2007; Graham *et al*., 2009). When functional and compositional turnover are paired with phylogenetic turnover, the latter informs us about the lability or conservatism of traits and the potential modes of speciation and biogeographical process underlying species distributions (Graham & Fine, 2008). Assuming that the environment plays an important role in determining the rate of species and functional turnover along an environmental gradient, a high phylogenetic turnover would be an indication that there is high niche conservatism that restricts close relatives to particular environments. In contrast, low phylogenetic turnover would be indicative of ecological speciation caused by local adaptation to different environmental conditions. In this case, the replacement of species along the gradient would happen mainly among close relatives, some of which may have originated in situ (Graham & Fine, 2008)

Even though combining the three metrics (i.e. functional, phylogenetic and compositional) of turnover provides a powerful test of niche versus stochastic processes (Graham & Fine, 2008), the structure of traits within local communities should further provide indications about the mechanisms underlying species turnover (McGill *et al*., 2006; Petchey & Gaston, 2002; Kraft *et al*., 2008, 2015). Species are a collection of traits that could evolve at different rates and respond differently to selective pressures (Ackerly & Cornwell, 2007). While some traits may vary stochastically as a product of genetic drift, other traits may vary deterministically according to different mechanisms (Ackerly & Cornwell, 2007). In addition, the scale at which selection operates is likely to vary among species. In plants, for example, traits such as rooting depth, leaf mass per area and the ability to fix nitrogen are traits that should respond locally to competition; alternatively, the degree to which a plant is deciduous and has compound or simple leaves is likely a response to climatic stressors such as drought and high temperatures (Kraft *et al*., 2015; Lebrija-Trejos *et al*., 2010). In birds, Miles & Ricklefs (1984) and Ricklefs (2012) suggested that overall morphology should respond to competitive interactions. Others have suggested that physiological tolerances of adults and juveniles should reflect adaptations to the environment (Webb, 1987; Kearney & Porter, 2009). Therefore, a combination of the measurement of trait turnover with the change in the structure of species traits along environmental gradients should allow us to not only differentiate between stochastic and deterministic community assembly, but could also reveal the mechanisms that determine community composition (McGill *et al*., 2006; Kraft *et al*., 2008, 2015).

In this study, we use compositional, functional and phylogenetic metrics of beta diversity to determine if the distribution of bird species along a steep environmental gradient in Colombia is deterministic or stochastic. Furthermore, we use ecological and morphological traits to test the hypothesis that the turnover in bird communities along the gradient is the product of a change in the mechanisms determining species composition along the gradient. Specifically, we predict that because dry forests are harsher and more stressful environments, the relative importance of species sorting through environmental filtering should be highest in dry forests. Thus, we expect that the trait space in physiological tolerances occupied by dry forest communities will be smaller than the one occupied by wet forest ones. In contrast, because of higher productivity and relaxed environmental filtering, wet forest communities should re spond more to biotic interactions such as competition and predation. In the particular case for competition, we expect wet forest communities to occupy broader eco-morphological trait space than dry forest communities.

## 2 Methods

### 2.1 Study Area

The Magdalena is one of the two lowland inter-Andean valleys occurring in central Colombia. The Magdalena River has been one of the most important rivers for navigation in the history of Colombia and of high importance in the colonization of northern South America. The river drops quickly from its headwaters to the lowlands in the upper Magdalena Valley, which is characterized by low annual rainfall (1000 mm). The low rainfall in the upper Magdalena is the product of the rain shadow of both the central and eastern Andes which rise above 4000 masl. About 200 km down river, the central Andes drop considerably in elevation allowing rainfall from the Pacific coast to pass over the Andes and fall in the mid Magdalena Valley increasing mean annual precipitation to almost 6000 mm in the western foothills of the eastern Andes. Because of its importance as a colonization route and as the connection for the interior of South America with the Caribbean Sea, the Magdalena Valley has a complex history of deforestation and fragmentation. The geological history of the Magdalena is also complex, because the wet forest has contracted and expanded several times during the last million years during glacial and interglacial periods. During the glacial periods, the entire valley was dry, which provided connections among the dry forest fauna and flora of the Caribbean region of Venezuela and Colombia and the dry forests in the upper Magdalena Valley (Haffer, 1967). During these periods, the wet forest fauna and flora were most likely restricted to refuges in the lowlands north of the Andes, Choco and southern Central America (Haffer, 1967).

### 2.2 Bird Sampling

We selected at random 15 localities along the Magdalena Valley distributed to capture the entire rainfall gradient (Table 1). In each locality we sampled birds using 50-m fixed radius point counts (Hutto *et al*., 1986) in which we counted all birds detected both visually and aurally for a period of ten minutes at each point. Point counts were repeated temporally a maximum of four times, although some of the points where only counted once (Table 1). Points were separated from the edge of the forest by a minimum distance of 75 m and were separated from each other by at least 200 m to ensure independence and minimize the sampling of species of the matrix surrounding the patch (Blake & Loiselle, 2001). Each morning we conducted up to ten point counts starting at dawn and until 10:00 AM or until activity dropped considerably. We avoided censusing during windy and rainy days. For the analyses we did not include Toucans, Parrots, Hummingbirds, Swallows, Swifts, water birds or birds that flew over the point while censusing because it was difficult to determine the independence of point counts for loud and highly mobile species. Because we did not have a large enough sample size to estimate the density of birds while correcting for detection probability, we estimated the abundance of bird species as the mean number of counts per species per point count.

**Table 1:** Location and description of localities sampled along the rainfall gradient of the Magdalena Valley. We show environmental variables as well as number of point counts and replicates per point count performed in each forest patch. Elev = Elevation (m), Bio12 = Annual Rainfall (mm), Dist = Distance to climatic station (Km), Bio2 = Mean Diurnal Temperature Range (° C), Bio5 = Max Temperature of Warmest Month (° C), Bio15 = Precipitation Seasonality (Coefficient of Variation), Bio17 = Precipitation of Driest Quarter (mm), Points = Number of census points for birds, Reps = Number of Replicates each point was censused

| Locality | Elev | Bio12 | Dist | Bio1 | Bio2 | Bio5 | Bio15 | Bio17 | Points | Reps |
| --- | --- | --- | --- | --- | --- | --- | --- | --- | --- | --- |
| Bateas | 429 | 1193.5 | 3.29 | 27.5 | 11.7 | 35.1 | 60 | 78 | 36 | 3 |
| El Triunfo | 196 | 1281.8 | 8.40 | 27.3 | 10.4 | 34 | 51 | 217 | 7 | 2 |
| Potosi | 400 | 1330.5 | 1.47 | 27.4 | 11.3 | 35 | 58 | 102 | 30 | 2 |
| ManaDulce | 490 | 1456 | 5.04 | 27 | 11.2 | 33.8 | 46 | 176 | 32 | 4 |
| Venadillo | 335 | 1599.7 | 10.39 | 27.5 | 11.3 | 34.5 | 45 | 174 | 20 | 2 |
| Jabiru | 341 | 1623.5 | 4.77 | 27.2 | 10.4 | 33.9 | 48 | 216 | 27 | 4 |
| Boqueron | 650 | 2249.6 | 6.37 | 26.2 | 10.8 | 32.6 | 44 | 219 | 13 | 1 |
| Mariquita | 475 | 2263 | 0.00 | 26.2 | 9.8 | 32.5 | 43 | 334 | 11 | 2 |
| Maceo | 639 | 2554 | 17.67 | 25.9 | 9.9 | 31.5 | 43 | 265 | 13 | 2 |
| Barbacoas | 138 | 2675.2 | 6.16 | 27.8 | 9.9 | 33.6 | 45 | 245 | 34 | 3 |
| Rio Manso | 160 | 2697.1 | 9.22 | 27.4 | 10.2 | 33.6 | 47 | 325 | 26 | 4 |
| La Perla | 300 | 2714.7 | 16.79 | 27.1 | 9.6 | 32.5 | 35 | 399 | 9 | 1 |
| San Juan | 168 | 2888.5 | 11.07 | 28.1 | 9.7 | 33.8 | 44 | 254 | 20 | 4 |
| Remedios | 718 | 2906.3 | 20.22 | 25.4 | 9.7 | 31.1 | 44 | 272 | 20 | 3 |
| Rio Claro | 449 | 3775.9 | 3.88 | 26.1 | 10.1 | 32 | 43 | 347 | 15 | 4 |

### 2.3 Community Turnover

We had two particular objectives in this study. The first was to test if there was a difference among compositional, functional and phylogenetic turnovers in relation to rainfall. In this analysis, we used three sources of information: abundance of species in each locality (as described above - mean number of individuals per point count), morphological and behavioral traits of each species and the phylogeny of all of the species we detected in our study. In the sections below, we provide detailed information about which traits we measured. For the phylogenetic comparisons, we downloaded 1000 trees, to account for phylogenetic uncertainty, from BirdTree.org using Hackett *et al*. (2008) as the back bone for the distribution of trees (see Jetz *et al*., 2012 for details on how the trees where constructed).

We calculated compositional turnover using the Chao index for assessing similarity of composition among communities while taking into account both abundance and sampling error (Chao *et al*., 2005). The Chao index is an extension of the Jaccard index, which incorporates a probabilistic framework to account for species abundances and the chance that species might be shared but, because of their rarity in either community, they might be considered as absent from one of the communities because of sampling limitations (Chao *et al*., 2005). The index estimates the probability that any two individuals sampled at random are shared by both communities while taking into account that shared species might be present in the communities but not sampled (Chao *et al*., 2005). Phylogenetic turnover was calculated as the total length of shared and unshared evolutionary history among any two communities denoted by the length of the branches in the phylogenetic tree shared among communities and unique to each community (Bryant *et al*., 2008). Because we had a distribution of phylogenetic trees we estimated phylogenetic turnover for each tree and report the mean turnover for the set of 1000 trees.

We calculated functional turnover in a similar way to phylogenetic turnover, but instead of using a molecular tree to determine relationships among species, we used trait data to infer a dendrogram of similarity among species (Petchey & Gaston, 2002). To construct the dendrogram, we used the total morphological matrix using both continuous and categorical traits. A detailed description of the traits and how they were measured is provided in the Trait Sampling section below. To allow categorical traits in the calculation of the dissimilarity matrix we used the general coefficient of similarity proposed by Gower (1971). Following the calculation of the dissimilarity matrix we performed a hierarchical clustering using the UPGMA method to construct the dendrogram, which performs better than other traditional methods in estimating species clustering for functional diversity analysis (Podani & Schmera, 2006). We performed calculations of compositional and functional turnover using vegan (Oksanen *et al*., 2016) package and phylogenetic turnover using the picante package (Kembel *et al*., 2010) in R (R Core Team, 2013).

We determined the relationship between community similarity and rainfall by performing a multidimensional scaling of the beta diversity and relating the first axis of the scaling to rainfall. This methodology allowed us to determine if the turnover happened linearly in relation to rainfall or had a logistic (stepwise) form, in which case we could estimate the amount of rainfall at which the community turnover is maximal. Additionally, using scaling of distance matrices or similar analyses such as canonical correspondence analysis provides stronger statistical power to detect the amount of community turnover that can be explained by the variation in the environmental gradient (Legendre *et al*., 2005). We compared the linear model with a logistic function model in which scaled community similarity was the dependent variable and rainfall the independent one. The logistic function was of the form

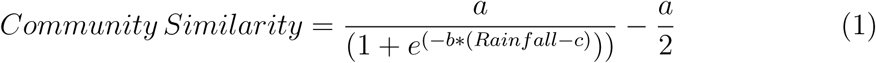

in which *a* determines the height of the curve and in this case the maximum difference estimated between types of communities, *b* determines how fast the transition happens from one type of community to another, and *c* determines the inflection point in which the community is expected to transition from type *x* to type *y*. To estimate the parameters of the logistic function, we used least squares minimization similar to a traditional linear regression. We then compared the models using Akaike Information Criterion (AIC) and *r*^2^. We performed the least squares minimization of the logistic function in R using the optim function.

Because the rainfall gradient of the Magdalena Valley expands over two ecoregions (Olson *et al*., 2001), we determined if the compositional, functional and phylogenetic turnover were higher than expected by chance between localities in different ecoregions and lower within ecoregion. In particular, if environmental filtering operates stronger in the dry forest than in wet forest we expected the turnover to be lower in localities in the dry forest than in localities in the wet forest indicating lower community variability. We constructed 1000 random communities using a swap algorithm that maintains the species abundance distributions as well as the richness of the communities (Hardy, 2008). For each of the 1000 random communities we calculated the compositional, functional and phylogenetic metrics using the observed functional dendrogram and phylogenetic tree. To determine if the turnover was higher or lower than expected by chance we calculated a standardized effect size (SES) for each of the metrics. The SES was computed as

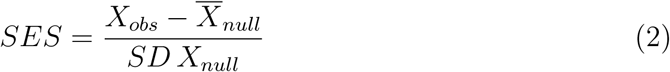

Overall, SES values higher than 1.96 or lower than -1.96 denote significantly higher or lower turnover than expected by chance, respectively. Finally, we determined if turnover within types of forests was lower than expected by chance using a t-test.

### 2.4 Climatic Description of Localities

To determine the influence of different environmental variables on the turnover of bird communities along the Magdalena Valley, we obtained mean annual rainfall and temperature variability from different sources. We obtained mean annual rainfall data from the closest climatic station to each of the localities sampled (IDEAM; Table 1). Climatic stations are run by Instituto de Hidrologia, Meteorologia y Estudios Ambientales (IDEAM) in Colombia. Mean rainfall for the period of 1981 - 2010 and the location of each station are freely available for download from their website. We determined the closest station to the locality by measuring geographic distance. Localities were all within 20 km of the closest station but most of them where much closer (mean = 8.31 km; Table 1). To account for possible deviations in rainfall due to distance from the station to the localities, we corroborated rainfall data from mean annual rainfall layer from bioclim (Hijmans *et al*., 2005). Because one way in which dry forests might be stressful to birds is through its stronger seasonality than wet forests, in addition to mean annual rainfall, we obtained information about rainfall seasonality and rainfall in the driest quarter from bioclim (Hijmans *et al*., 2005). Mean annual temperature, mean maximum temperature and temperature range were also obtained from bioclim (Hijmans *et al*., 2005). Finally, to obtain temperature variability, we used five Hobo U23 data loggers that were located in two dry forests and three wet forests. The data loggers were set to measure temperature and relative humidity each hour for an average of 662 days (Mana Dulce = 585 days, Jabiru = 233 days, Rio Manso = 730 days, San Juan = 659 days and Rio Claro =1104 days). Finally, we tested for significant differences in mean annual temperature, temperature range, mean maximum temperature, temperature coefficient of variation, precipitation seasonality and precipitation in driest month using a linear model relating each of this variables to rainfall in each locality.

### 2.5 Trait Sampling

In order to determine the influence of different mechanisms that could determine the rate of community turnover along the gradient, we constructed a database with morphological and ecological traits hypothesized to vary according to environmental filtering and competition. Below, we will describe the traits and predictions of how we expect the morphological trait space to vary depending on the mechanisms hypothesized to operate in each locality.

#### 2.5.1 Environmental Filtering

To test the hypothesis that the relative importance of environmental filtering is higher in dry forests, we obtained data on species’ climatic preferences. We specifically wanted to test the hypothesis that species that occupy the dry forests in the Magdalena Valley experience more stressful conditions throughout their ranges. By stressful conditions we mean explicitly higher maximum temperatures, wider temperature ranges, higher precipitation seasonality and lower precipitation during the dry seasons. All four variables potentially affect species distributions directly or indirectly. For example, wider temperature ranges and higher maximum temperatures might be problematic for the eggs (Webb, 1987) and potentially the adults (McKechnie & Wolf, 2009). Additionally, high precipitation seasonality and low precipitation during dry season are problematic for water regulation in both adults and nests but also might affect species through resource availability, which might be much lower during the dry season for most (but not all e.g. nectarivores) foraging guilds. To test the environmental filtering hypothesis, we obtained the mean values of diurnal temperature range (bio2), maximum temperature in warmest month (bio5), precipitation seasonality (bio15) and precipitation of driest quarter (bio17) for each species throughout their range. We then computed community-wide environmental tolerances by computing the mean of the species present in the community weighted by the abundance of each species. For the former analysis we assumed that climatic variables are a good proxy for environmental stressors for species and thus for their physiological tolerances. Environmental data were obtained from bioclim (Hijmans *et al*., 2005) and species distribution ranges from bird life international database (Birdlife International & NatureServe, 2014). Another prediction of the environmental filtering hypothesis is that in communities with stressful environments the species should have more similar environmental tolerances among them than species in more benign environments. Therefore, communities in dry forests should occupy a smaller trait space than wet forest communities. To test this prediction, we estimated functional richness and dispersion using community-wide measurements of temperature range, maximum temperature, rainfall seasonality and minimum rainfall. Functional richness is defined as the volume of the convex hull polygon delimited by the values of the n traits and s species present in the community (Cornwell *et al*., 2006; Villéger *et al*., 2008). Functional Dispersion estimates the morphological centroid of the community in response to species abundances, and then estimates the spread of species from the centroid of the community (Laliberté & Legendre, 2010). Finally, using the same randomization procedure described previously to test for significance in the turnover of communities, we constructed 1000 random communities, and calculated a SES to determine if functional richness and dispersion of physiological tolerances were smaller than expected by chance particularly in dry forests.

Additionally, we sought to test the hypothesis that dry forest species better regulate the temperature of their nests than wet forest birds. Specifically, we wanted to determine if dry forest birds had greater differences between maximum internal and maximum external temperatures throughout their development to determine if selection to avoid temperature extremes may be stronger in dry forests. Also, we explored if dry forest nests had lower internal temperature variability relative to the ambient temperature variability. We followed the development of 57 nests from 23 species in two localities, one in dry forest and one in wet forest (Dome = 6, Cup = 45, Platform = 6). We recorded temperature inside and 10 cm outside of the nest with a Hobo USB data logger for the length of the entire development of the nest or until it was either abandoned, depredated or nestlings died. We then calculated the difference between the maximum ambient and nest temperatures and the ratio between inside and outside temperature variance. As the ratio converges on one, outside and inside temperature variance are similar. If the ratio is less than one, it means that nest temperature variability is lower than ambient temperature variance and thus the nests are able to dampen environmental variability. Thus, nests that are adapted to avoid extreme changes in temperature that would affect the nest should show a larger difference between inside and outside temperature and a lower than one temperature variability ratio. We asked if there were differences in temperature regulation within nest types among forest types using two sampled t-test. The p-value of all of the t-tests performed were adjusted using bonferroni test correction.

#### 2.5.2 Biotic interactions: Competition

To test the competition hypothesis, which predicts that competitively structured communities will be more overdispersed in morphological trait space, we measured eight morphological traits and one ecological trait that have been suggested to be correlated with the ecology of species (Pigot *et al*., 2016; Ricklefs, 2012; Miles & Ricklefs, 1984). The traits were: body mass, wing length, tail length, bill width, depth and height and tarsus length and foraging strata. The morphological traits were collected in the field using mistnets to capture birds. Tarsus and bill measurements were collected using a caliper with 0.01 precision and tail and wing length were measured using a wing ruler. Morphological data were available for 123 of the 213 species accounted in the analysis. For most of the species we used the mean of at least two individuals but some were represented by only one specimen (n=1225 individuals, mean number of individuals per species = 10, Number of species with two or more individuals = 92, Number of species represented by one individual = 31). We are aware that the species represented by one individual can potentially bias our analysis, but it is likely that in all of the species within the Magdalena Valley, intraspecific trait variation is smaller than interspecific variation. We estimated body mass for the species that we did not have morphological measurements using the CRC hand book of avian body masses (Dunning Jr, 1992). For foraging strata, we used a recently published database for all the bird species of the world (Wilman *et al*., 2014). The database separates foraging strata into five separate categories; ground, understory, mid story, canopy and aerial and for each category assigns a proportion of time that the species spends in each stratum. In that way, foraging stratum can be treated as a quantitative trait instead of a categorical one. To maintain foraging stratum as a single trait we coded the stratum from 1 - 5 sequentially from the ground to aerial foraging. Subsequently, we obtained the weighted average for each species, with weights determined by the percent use of each strata. For example, if a species forages 20% of the time in the ground, 40% in the understory and 40% in the mid story the foraging strata value for that particular species was calculated as *Foraging Stratum* = 0.2 × 1 + 0.4 × 2 + 0.4 × 3 + 0 × 4 + 0 × 5 = 2.2.

Using the nine eco-morphological traits, we performed a principal components analysis (PCA) to reduce collinearity among variables. Because the variables ranged over several orders of magnitude, all of the variables were centered to have a mean of zero and scaled to have variance of one prior to the PCA analysis. We used the rotated scores from the first five PCs (the first five components explained 99.5% of the variation) to calculate functional richness and dispersion indices (i.e. eco-morphological richness and evenness). In competitively structured communities, the prediction is that ecomorphological richness and dispersion are higher than in communities structured by environmental filtering. Given the previous definitions of the metrics of functional diversity, the competition hypothesis predicts that there should be an increase in both functional richness and dispersion with rainfall along the gradient (Pigot *et al*., 2016; Kraft *et al*., 2008). The significance of the relationship between eco-morphological richness and rainfall was assessed using a least squares linear regression.

Additionally, to determine if eco-morphological richness and dispersion were higher in wet forest than expected by chance, we constructed 1000 random communities for each of the 15 localities using the entire source pool of the Magdalena Valley, but maintaining both the frequency of each species and the richness of communities. To assemble the random communities we used the independent swap algorithm over 1000 iterations (Gotelli, 2000; Hardy, 2008). For each of the 15000 communities, we then calculated eco-morphological richness and dispersion. Finally, we calculated a SES richness and dispersion for each community with the expectation that wet forest communities would have SES values of richness and evenness greater than 1.96.

In addition to the competition for food resources, species might also compete for nest space (Martin, 1988). This hypothesis predicts that species in the wet forest would have more diversified nesting strategies than in dry forests. To test the latter prediction, we calculated nest diversity and dispersion among communities in a similar way than for eco-morphological traits. In this case, because the trait is categorical, functional richness was measured as the number of unique trait combinations in the locality (i.e. number of nest types, Villéger *et al*., 2008). Functional dispersion was calculated in a similar way as described above. Because in the case of nest richness the data are counts of species with the same type of nest, we tested the significance of the functional richness and rainfall relationship using Poisson regressions. The relationship between nesting dispersion and rainfall was assessed using a beta regression. Functional dispersion and richness metrics were calculated using the FD package (Laliberté & Legendre, 2010; Laliberté *et al*., 2014) in R.

## 3 Results

### 3.1 Compositional, Functional and Phylogenetic Turnover

We found support for a stepwise turnover pattern in composition, function and phylogeny. In all of the cases, the logistic model fit the data better than a simple linear model even though it had at least one more parameter (Table 2, Figure 1). Rainfall explained on average 88% of the variance in community similarity along the gradient (Table 2). The maximum turnover of the communities occured around the 2300 - 2400 mm isocline consistently for the three types of turnover measurements. Communities above and below the 2300 mm isocline are on average 75% different according to the compositional turnover, 64% to the functional turnover and 58% to the phylogenetic.

**Table 2:**
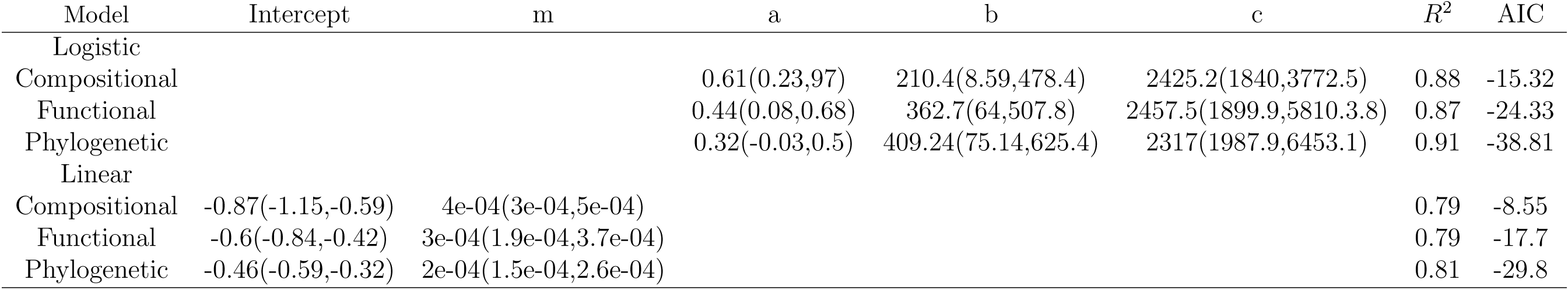
Results of model selection for the relationship between community similarity and rainfall. Intercept and m are the parameters of the linear model and a, b and c are the parameters estimated for the logistic model. The best model is the one with the lowest AIC and significant differences among models are detemined by differences greater than 2 in their AIC.

**FIGURE 1:**
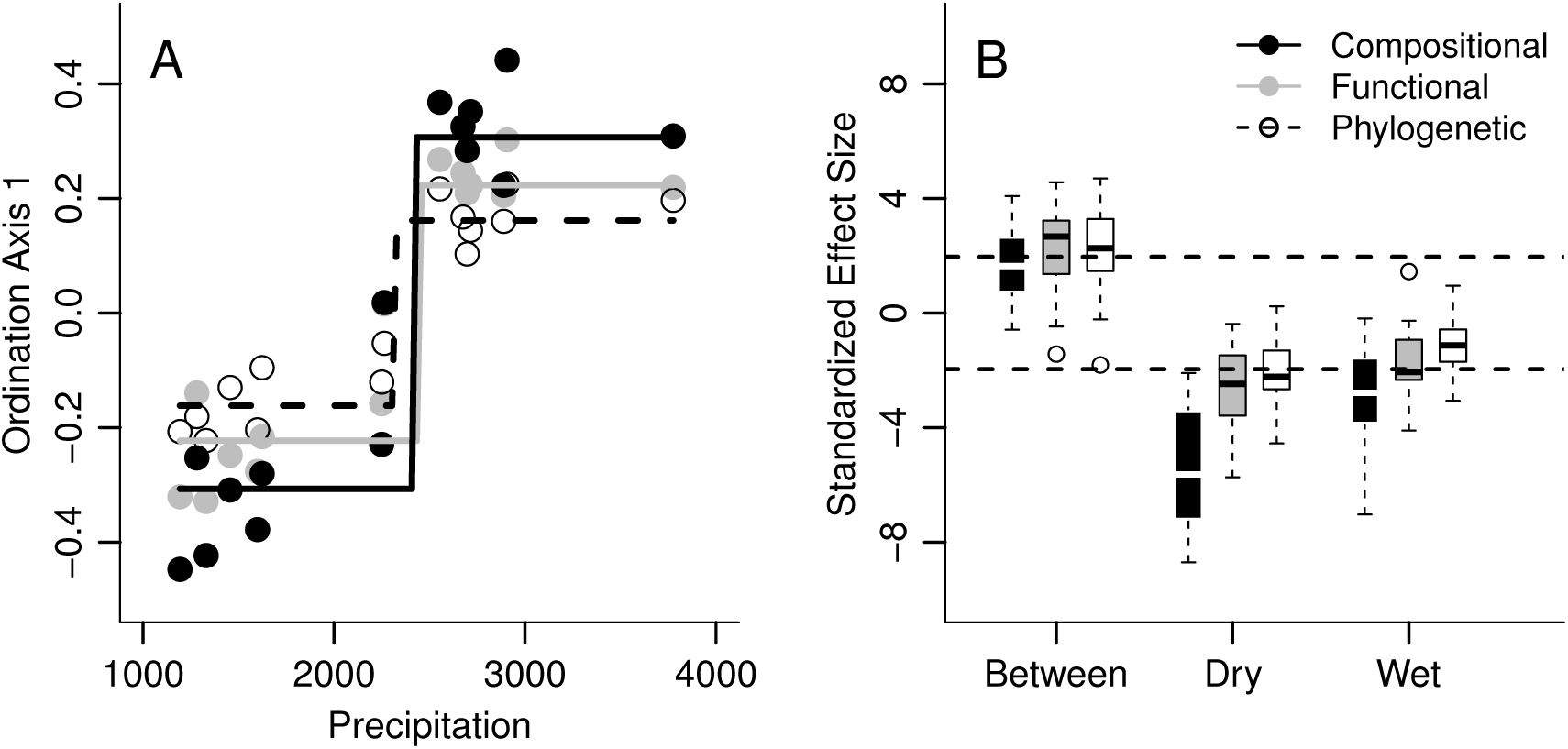
Compositional, functional and phylogenetic turnover of lowland bird communities along the rainfall gradient of the Magdalena Valley, showing (A) a steep turnover around the 2300 mm rainfall isocline that is consistent among the measurements, but the measurements decrease in strength of turnover from compositional to phylogenetic, and (B) shows the distribution of the Standardized Effect Sizes for three types of comparisons: between and within wet and dry forests, showing higher than expected by chance turnover between types of forest and lower than expected by chance turnover within dry forests.

Thirty-seven percent, 71% and 61% of the communities had higher compositional, functional and phylogenetic turnover, respectively, than expected by chance between types of forests (Figure 1). Within the dry forest all of the comparisons where smaller than expected by chance according to the compositional turnover. Functional and phylogenetic turnover showed that 71% and 67% of the comparisons respectively had smaller turnover than expected by chance, respectively. Within wet forest, on the other hand, the rates of compositional, functional and phylogenetic turnover showed that 67%, 52% and 19% of the comparisons where significantly smaller than expected by chance. Finally, the dry forests were significantly less variable than the wet forests, as suggested by a lower mean compositional, functional and phylogenetic SES (Compositional; mean dry = -5.4, mean wet = -2.92, t = 3.9, df = 38.8, p>0.01; Functional; mean dry = -2.6, mean wet = -1.6, t = -3.5, df = 38.5, p=0.001; Phylogenetic; mean dry = -2.19, mean wet = -1.1, t = 3.29, df = 39.2, p>0.01).

### 3.2 Environmental Variables

We found that while mean annual temperature was constant among localities (*Temperature* = 27.7-3.8 × 10^−4^ × *Rainfall*; *p* = 0.16), both temperature range (*Temperature Range* = 11.9 - 6.8 × 10^−4^ × *Rainfall*; *p* < 0.01, *r*^2^ = 0.57) and mean maximum temperature (*Max Temperature* = 35.9 - 1.1 × 10^−3^ × *Rainfall*; *p* < 0.01, *r*^2^ = 0.53) decreased with rainfall. Also, the coefficient of variation of hourly temperature decreased significantly with rainfall (*Temperature CV* = 18.19-3 × 10^−3^ × *Rainfall*; *p* < 0.01, *r*^2^ = 0.93), suggesting that temperature is significantly less variable as rainfall increases. Finally, rainfall seasonality and rainfall in the driest month significantly increased along the gradient (*Seasonality* = 58.4 - 5 × 10^−3^ × *Rainfall, p* < 0.01; *r*^2^ = 0.45; *Min Rainfall* = 39.7 + 0.09 × *Rainfall, p* < 0.01; *r*^2^ = 0.62)

### 3.3 Environmental Filtering

We found that community temperature range and rainfall seasonality decreased significantly (*Temperature Range* = 15.05 + *Rainfall*^(-0.054)^; *r*^2^ = 0.59; *Rainfall Seasonality* = 68 - 4.6^−3^ × *Rainfall*; *p* < 0.01; *r*^2^ = 0.85) and community minimum rainfall to increase with annual rainfall in each locality (*Minimum Rainfall* = 66.3 + 0.05 × *Rainfall, p* < 0.01; *r*^2^ = 0.82; Figure 2). We found no relationship between rainfall and community maximum temperature (*Maximum Temperature* = 31.9-1 × 10^−4^× *Rainfall, p* = 13; *r*^2^ = 0.15; Figure 2). Physiological trait structure did not follow our predictions. The trait space in physiological tolerances was not smaller or less dispersed in dry forests as expected. This is shown by the non-significant relationship between community physiological richness or dispersion and rainfall (*Richness* = 20.4+6.8 × 10^−4^ × *Rainfall, p* = 0.8; *Dispersion* = 1.44+6.5 × 10^−6^ × *Rainfall, p* = 0.9). Finally, neither physiological richness nor dispersion of dry forest communities was lower than expected by chance.

**FIGURE 2:**
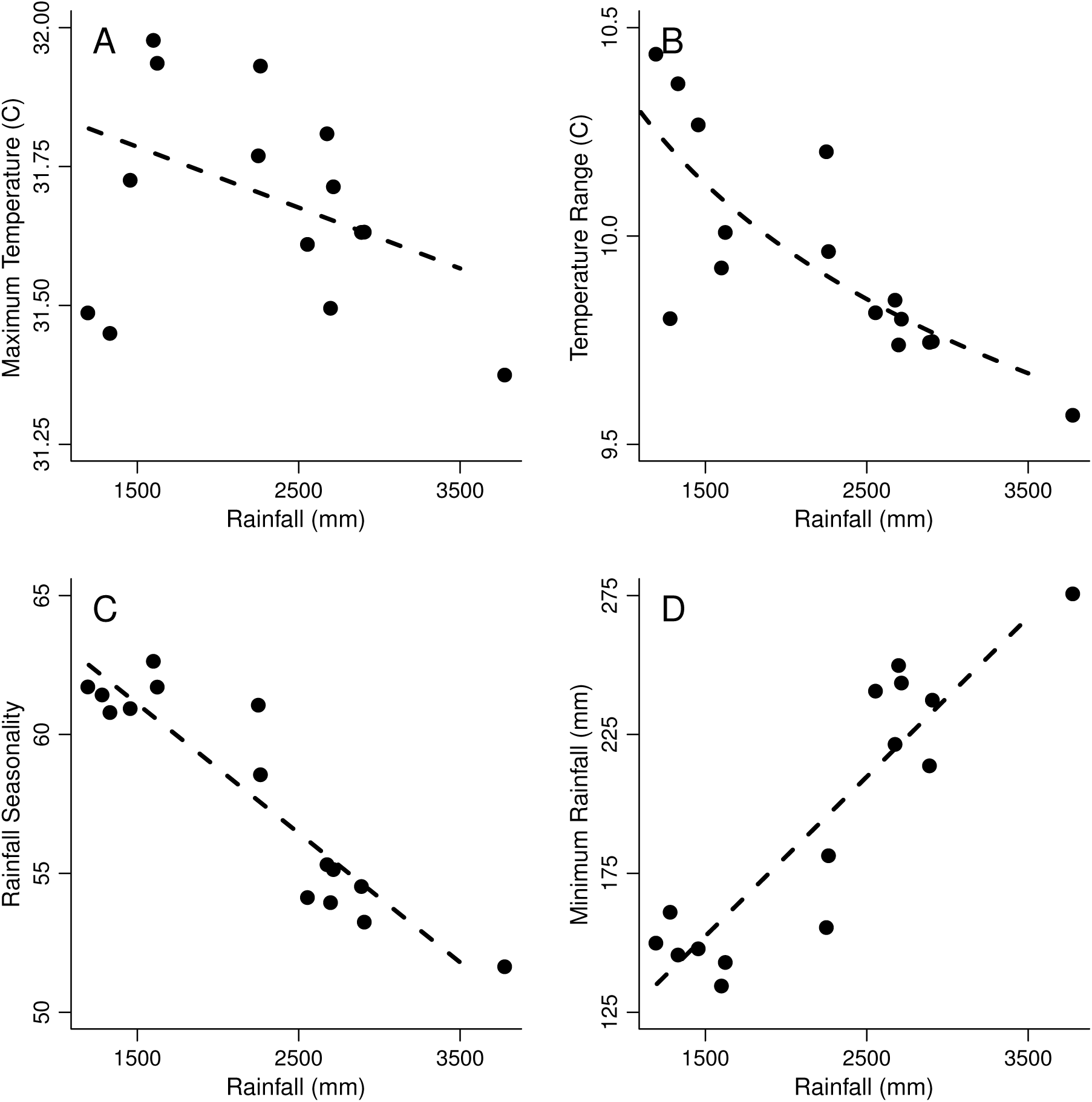
Relationship between average community physiological tolerances and rainfall, showing, (A) no relationship between community average maximum temperature tolerance; (B) a negative relationship between average community temperature range and rainfall; (C), a decrease in mean community rainfall seasonality with locality rainfall and; (D) an increase in the minimum rainfall that species experience throughout their ranges.

Among nest types, we found that cup and platform nests in dry forests had significantly lower differences between internal and external max temperatures. Only cup nests in dry forests had significantly lower internal variance relative to the environmental variance when compared to wet forests (Table 3). In fact, the temperature variance in cup nests of dry forests was significantly lower than ambient temperature (*Mean* = 0.5, *df* = 3, *p* = 0.03). This result means that variance in temperature of cup nests in dry forests was 50% lower than the variance in ambient temperature. The variance in nest temperature of platform nests in dry forests was also 45% lower than ambient variance, but this difference was not significant (*Mean* = 0.55, *df* = 1, *p* = 0.06)

**Table 3:** Results of t-tests comparing max difference and variance ratio between nest and ambient temperature among nests in dry and wet forests. The results are product of multiple t-tests comparing types of nests and each type of nest between localities. For comparison among types of nests the objective was to determine if the difference between nest and ambient temperature was less than zero and less than one in the case of the ratio of variances.

| Type of Nest | Dry | Wet | df | p |
| --- | --- | --- | --- | --- |
| Max Difference |  |  |  |  |
| Cup | 4.69 | 9.6 | 8.5 | > 0.01 |
| Dome | 5.93 | 4.17 | 2.96 | 1 |
| Platform | 3.65 | 9.12 | 2.99 | 0.02 |
| Variance Ratio |  |  |  |  |
| Cup | 0.5 | 3.57 | 36.5 | > 0.01 |
| Dome | 1.63 | 1.87 | 3.71 | 1 |
| Platform | 0.55 | 1.46 | 2.07 | 1 |

### 3.4 Biotic Interactions: Competition

As rainfall increases, the strength of environmental filtering should decrease and thus competition for resources should be more important in determining community structure. The competition hypothesis predicts that species co-occurring locally should be ecologically and consequently morphologically more diverse to avoid competition (MacArthur & Levins, 1967), but we found no evidence for change in eco-morphological richness or dispersion with increasing rainfall (*Functional Richness* = 86.7 - 2 × 10^−4^ × *Rainfall, p* = 0.99; *Functional Dispersion* = 1.45 - 9.8 × 10^−6^ × *Rainfall, p* = 0.9). Furthermore, only one site in the wet forest (Barbacoas) had eco-morphological richness higher than expected by chance (Figure 3). We found no relationship between nest richness and rainfall (*Nest Richness* = 5.25+3.9^−4^ *Rainfall, p* = 0.26), but nest dispersion increased with rainfall as predicted (*Nest Dispersion* = 0.23 + 2.99 × 10^−5^ *Rainfall, p* < 0.01; *r*^2^ = 0.58).

**FIGURE 3:**
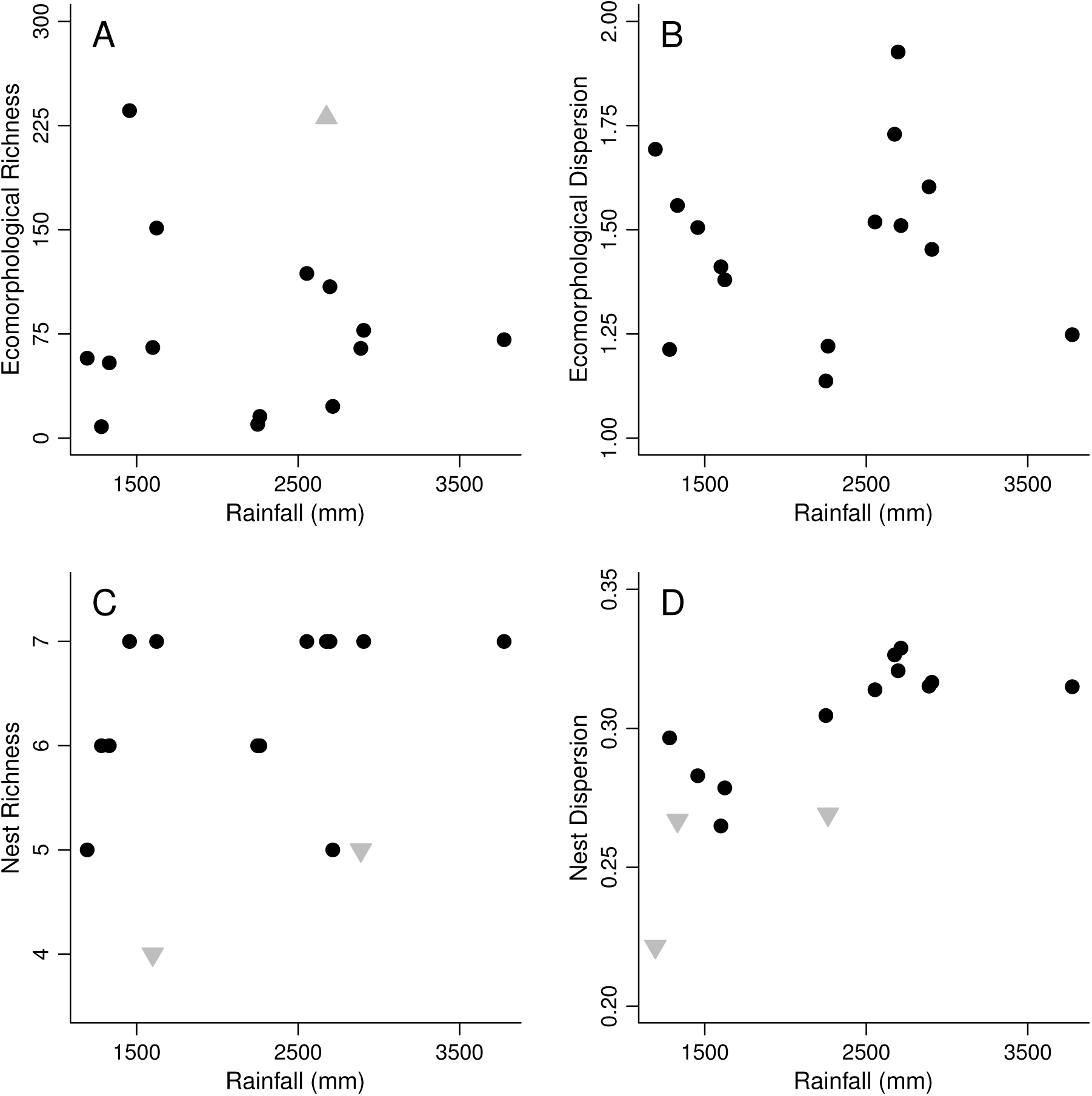
Eco-morphological and nest structure of communities along the rainfall gradient of the Magdalena Valley. A and B show no relationship between ecomorphological richness and dispersion and rainfall. C shows a slight but not significant increase in nest richness with rainfall and D shows a significant increase in nest dispersion throughout the gradient. Grey triangles indicate localities in which functional richness or dispersion was higher (triangles pointing up) or lower (triangles pointing down) than expected by chance

## 4 Discussion

Our results suggest that there is deterministic bird community turnover around the 2400 mm annual rainfall isocline in the Magdalena Valley in Colombia. The rainfall gradient promoted a strong compositional, morphological and phylogenetic turnover in which almost the entire community was replaced in a short geographic distance. The models are strongly consistent with a stepwise function replacement of the communities (Table 1). Around the 2400 mm isocline there is up to a 75% change in the community, whereas in more than 200 km of dry or wet forest spanning a rainfall gradient of more than 1000mm on either side of this transitional zone, the average turnover among communities within the same type of forest was only 41%. Furthermore, our results partially suggest that environmental filtering might be of higher importance for structuring communities in the dry end of the Valley. Not only did dry forest communities have significantly less turnover than expected by chance, they also had lower turnover rates than wet forest communities (Figure 1). Species in the dry forests were also better adapted to higher rainfall seasonality and stronger dry seasons (Figure 2). In wet forests, we found evidence that competition for nest sites is stronger than in dry forests and the lower phylogenetic turnover compared with compositional turnover might be an indication of replacements among closely related ecologically similar species that do not coexist because of competition (Robinson & Terborgh, 1995). Nevertheless, there was little evidence that the communities were more dispersed in traits in wet forests than in dry forests suggesting that competitively structured niches are not necessarily more likely in wet than in dry forests.

Differences in temperature regulation within nest types between types of forests also point to the possibility that climate might be a determinant of community composition in dry forests. Cup and platform nests in the dry forests dampen the high environmental variability of the habitat whereas they do not in wet forests (Table 3). Our data also show that the difference in maximum inner and outer nest temperature is lower in cup and platform nests of dry forests, suggesting that that species might be more selective of the microclimates of nest sites in the dry than in the wet forests (Table 3). Such patterns might also result from higher nesting seasonality in dry forest birds, which may only nest during the wet season when temperature variation is less extreme. Thus, an alternative prediction that arises from the environmental filtering hypothesis is that there should be a decrease in nesting seasonality with increasing rainfall. Some studies suggest that in Amazonian wet forests birds nest throughout the year, ignoring rainfall seasonality (Stouffer *et al*., 2013). In dry forests, however, we have no comparable data on the nesting phenology that could potentially support our hypothesis and predictions.

One caveat that rises against the environmental filtering hypothesis is the low support for the prediction that dry forest species should be exposed to higher temperatures throughout their ranges and that physiological trait space in dry forest communities should be smaller and less dispersed compared to wet forest communities (Figure 2). On average species in the dry forests are not exposed to higher temperatures throughout their ranges than wet forest species and there was no relationship between physiological trait richness and dispersion and rainfall. Also, even though the relationship between mean temperature range of species in the community and rainfall differed significantly, the magnitude of the decrease was less than in 0.5° C, which might not necessarily represent a strong selective agent. It is possible that the resolution of the environmental layers used to collect the data throughout the ranges of the species was not high enough to capture the real strength of the environmental filtering in dry forest. First, our data logger captured hourly and daily variability that were not represented in the broad-scale data. The data obtained across the ranges of species were rough estimates of mean maximum annual temperature and monthly temperature ranges. The hypothesis specifically deals with daily temperature in a few hours in a portion of the days of the year were temperature rises above 40° C.

Birds can potentially compete for nest resources, which might influence community assembly (Martin, 1988). We found support for this hypothesis as the dispersion in nesting types increased significantly with rainfall. Such patterns further support a shift in the mechanisms that drive community composition along the gradient. One of the ways that environmental filtering may be operating in the dry forests is through high variability and extreme high temperatures in the dry forests. Such mechanisms would predict lower functional dispersion of nesting types as the nests that better regulate temperature should be selectively favored in this type of forest. We provide some evidence that cup and platform nests in dry forests better regulate temperature than the same types of nest in wet forests, in which temperature extremes may not be great enough to require regulation of the microclimate. Nevertheless, our results indicate that temperature is a potential determinant of species composition and/or behavior. The increase in rainfall was associated with a decrease in temperature variability and maximum temperature. If temperature regulation is not a problem in wet forests, it opens the possibility of a diversification in nest types to decrease the impact of competition. In dry forests, however, the reduction in nest types could increase competition as it is more critical for species to select for the best places to locate nests and avoid high temperatures. Thus, environmental filtering may increase competition for a potentially limiting resource (i.e., nest sites), which could further constrain which species can occur in a community.

Our functional trait data do not support the hypothesis that there is stronger competition for resources in the wet forests as there was no difference in the trait space of wet and dry forests. Alternatively, competition for resources in the dry forest may occur at similar levels in both communities. Many studies have inferred that competition is an important determinant of bird species distributions and abundance (Jankowski *et al*., 2012), but few of these studies were conducted in dry forests, which have been historically understudied (Oswald *et al*., 2016). Thus, our data suggest that in addition to the environmental filtering, competition for resources might also influence dry forest communities. However, neither eco-morphological richness nor dispersion was higher than expected by chance in any of the localities. The other potential explanation is that the morphological traits are not related to the niche axes that experience competition (Miles & Ricklefs, 1984; Ricklefs, 2012) or that the relationship between ecology and morphology is much more complex than previously thought (Pigot *et al*., 2016). Thus, it is also possible that that competition happens through other niche axes that we were unable to detect in this study.

One hypothesis that remains to be tested is the possibility that predation is stronger in wet forests, influencing community assembly (Martin, 1988; Jankowski *et al*., 2012). Many of the most important nest predators were only found in the wet forest. Preliminary data suggest that the three toucan species exclusively found in wet forest during my study (unpublished Data) are strong nest predators in these forests of the Magdalena Valley (G. Londono, unpublished data). In addition to the toucans, the number of forest raptors also increases as well as the richness of primates (Gomez et al unpublished data) in wet forests. While it has been hypothesized that cavity nests might protect the eggs and nestlings from heavy rainfall (Oniki, 1985), there is more evidence to suggest that this type of nest provides protection against predators (Oniki, 1979, 1985). Thus, increased predation pressure in wet forests might select for the observed increase in cavity nesters and a decrease in cup nesters with rainfall. Our data support this prediction (Figure 4) but the main assumption – that nest predation in dry forests is significantly lower than in wet forest – remains to be tested. Thus, our data are inconclusive about this hypothesis which we believe might be an interesting one to test in the future.

**FIGURE 4:**
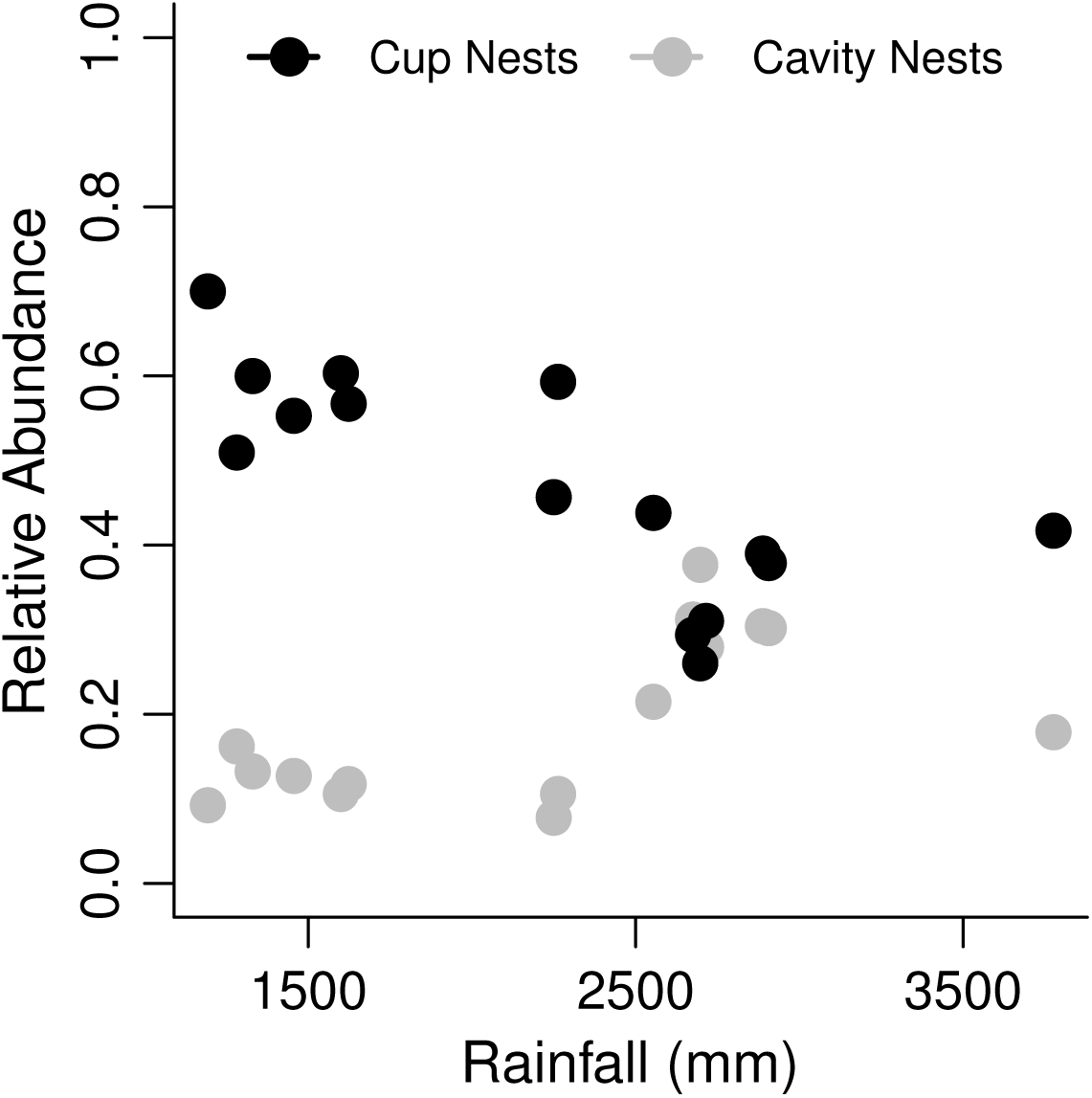
Relative abundance of Cavity and Cup nests along the rainfall gradient of the Magdalena Valley. Gray and black dotted lines are the fitted lines from a beta regression for the relationship between cup and cavity nests and rainfall.

In our study functional and phylogenetic turnover were lower than compositional turnover. A lower functional than compositional turnover suggests that there are some similar niches to be filled in both wet and dry forests, even though the niches are filled by different species with the same functional traits. This scenario would support a turnover mediated by interspecific competition (Robinson & Terborgh, 1995). In their work, Robinson & Terborgh (1995) report that intrageneric replacements along a productivity gradient in lowland Amazonia responded to inter-specific aggression between ecologically similar species. The heavier congener almost always actively displaced the smaller congener from the sites with higher productivity. We found several examples of replacement among ecologically similar species along the Magdalena Valley that fit this scenario. For example, white-bellied antbird (*Myrmeciza longipes*) in the dry forest is replaced by the chestnut-backed antbird (*Myrmeciza exsul*) in the wet forest. Both forage in similar habitats, close to the ground and potentially searching for similar insect items. An other example is the replacement of the endemic Apical flycatcher (*Myiarchus apicalis*) with its close relative dusky-capped flycatcher (*Myiarchus tuberculifer*). Both of this examples as well as some other ones occur among close relatives most likely in the same genus. Such patterns would lead to lower phylogenetic turnover. The functional and phylogenetic turnover, however, are still higher than expected by chance between forests and lower than expected by chance within forests suggesting a high change in function and evolutionary history of these communities with rainfall (Figure 1).

Even though we found a mismatch in the amount of turnover among compositional, functional and phylogenetic metrics, there is a spatial congruence in where the turnover happens (Table 2, Figure 1). The three metrics predict that the community shift happens at the boundary delimiting the Magdalena dry forests and Magdalena-Uraba moist forests ecoregions (Olson *et al*., 2001). Provided that the ecoregions of northern South America where delimited by vegetation data (Olson *et al*., 2001), this suggests a spatial match in the turnover of bird and plant communities along the rainfall gradient. Others have found strong associations between the turnover of plant and bird communities (Jankowski *et al*., 2013), suggesting that vegetation might have a very strong influence on the structuring tropical bird communities. There might be direct and indirect effects of vegetation on bird communities but we hypothesize that in the case of the Magdalena Valley the effects are direct. The dry forest tree community is mainly deciduous, such that in the dry season, the entire forest loses its canopy over, potentially increasing temperatures inside the forest, at least during the day. In the wet forest, the canopy is more permanent throughout the year, which stabilizes temperature and eliminates the strong filtering by high temperatures. This hypothesis predicts that the limits of the dry and wet forest are associated with a strong change in the proportion of deciduous trees that compose the canopy.

In conclusion, we provide evidence that suggests that the mechanisms driving community assembly along the Magdalena Valley in Colombia change with precipitation. In localities with low rainfall (<2400 mm), we found evidence for environmental filtering, whereas in localities above the 2400 mm isocline we found only partial evidence supporting stronger biotic interactions (e. g., predation and nest site use). This change in mechanisms can potentially explain the strong compositional, functional and phylogenetic turnover that happens abruptly over a short geographic distance. The Magdalena River has been one of the major centers for development in Colombia since colonial times. The high within-forest community variability might reflect this long history of fragmentation and deforestation (Harrison, 1997; Pardini *et al*., 2005). We report here that the Magdalena Valley bird communities might be two separate entities with high functional and phylogenetic diversity. Despite its high diversity and high levels of fragmentation and deforestation, there are no protected areas in the region. Our data suggest that the upper and middle Magdalena Valley must be treated separately in conservation strategies.

## 5 Acknowledgements

We thank Hda. Los Limones, C. Garcia, C. Mendoza, H. Llara, Remanso del Sumapaz, Pizano-Gomez Family, A.M. Jaramillo, Rio Claro, J.L. Toro, Corantioquia, A. Link, O. Laverde and G. Campuzano for support in the study sites. J. Drucker, A. Echeverri, J. Llano-Mejia, M.A. Loaiza, A. Morales-Rozo, E. Ortiz-Acevedo, J.L. Parra, J. Sandoval and E. Yepes for their assistance in the field. Funding sources Chapman AMNH, COS, Sigma-Xi and National Geographic-Waitts Grant No. W270-13 to J.P Gomez and NSF grant DEB 213858 to S.K. Robinson.

